# Memory for non-painful auditory items is influenced by whether they are experienced in a context involving painful electrical stimulation

**DOI:** 10.1101/341891

**Authors:** Keith M. Vogt, Caroline M. Norton, Lauren E. Speer, Joshua J. Tremel, James W. Ibinson, Lynne M. Reder, Julie A. Fiez

## Abstract

In this study, we sought to examine the effect of pain on memory. Subjects heard a series of words and made categorization decisions in two different contexts. One context included painful shocks administered just after presentation of some of the words; the other context involved no shocks. For the context that included painful stimulations, every other word was followed by a shock and subjects were informed to expect this pattern. Word lists were repeated three times within each context in randomized order, with different category judgments but consistent pain-word pairings. After a brief delay, recognition memory was assessed. Non-pain words from the pain context were less strongly encoded than non-pain words from the completely pain-free context. An important accompanying finding is that response times to repeated experimental items were slower for non-pain words from the pain context, compared to non-pain words from the completely pain-free context. This demonstrates that the effect of pain on memory may generalize to non-pain items experienced in the same experimental context.

## Introduction

The experience of acute somatic pain is often memorable. Aside from the memory of pain itself, noxious stimuli can affect learning and memory in several important ways. Brief painful experiences can enhance learning. After touching a hot stove, for example, an individual typically forms strong memories of the experience and learns to avoid this behavior in the future. However, the relationship between pain and memory is nuanced, and everyday examples of pain *impairing* memory are also common. Protracted pain can interfere with concentration and the ability to recall specific details of events that occur, for example, during a severe headache. Thus, the effect of pain on memory may depend on the context in which encoding occurs. Despite how commonplace such experiences are, empirical evidence quantifying pain’s effect on memory is limited.

The effect of pain on human memory has been interrogated using explicit memory testing, often for viewed images of scenes or objects. As a characteristic example, when tested several minutes later in a surprise recognition task, memory was worse for pictures paired with painful thermal stimulation, compared to those not paired with pain (Forkmann et al. 2016). In a similar image-categorization paradigm, association with painful electric shocks had no effect on immediate recognition memory testing, suggesting no effect for pain on memory (Schwarze et al. 2012). However, in a second cohort who underwent the same encoding experience but who were tested the following day, memory performance was enhanced for pain-paired items (Schwarze et al. 2012). These disparate results have several possible explanations, including the obvious experimental design differences. In addition to different intervals between encoding and memory testing, the type, duration, consistency of pairing, and timing of the pain stimulus vary across experiments, all of which may contribute to disparate results. These subtle design differences pervade the literature that examines pain’s effect on explicit memory and highlight why more confirmatory work is needed to discern which factors significantly modulate the interaction of pain on during memory formation. In this experiment, we tested for recollection and familiarity to auditory items, shortly after the encoding opportunity.

The present study was designed to examine two effects of interest: the effect of pain on memory for individual experimental items, and the effect of pain context on memory for non-pain items. Our hypotheses were designed around these two fundamental comparisons. For explicit memory, we first predicted that pain would impair memory performance on an item-level, within the pain portion of the experiment. Specifically, pain-paired words would show lower recognition compared to non-pain items interleaved in the same experimental context. Second, we anticipated that pain context would enhance memory performance due to increased overall arousal during that condition. Thus, we predicted that nonpain words interleaved with pain-paired words would be more strongly encoded than non-pain words experienced in the portion of the experiment described to subjects as being free from painful stimulation.

Not all memories can be explicitly recalled, and acute pain is known to also affect implicit memory in humans, mostly explored in fear conditioning research. These experiments demonstrate that, for items previously associated with noxious stimuli, re-exposure to the conditioned stimulus causes a sympathetic response, which can be measured by changes in heart rate (Castegnetti et al. 2016) or electrodermal activity (Schiller et al. 2010). Interestingly, the sympathetic response from this learned association may persist up to one year later (Schiller et al. 2010), indicating the possibility of a prolonged aversive association for items previously experienced with pain. An interesting subset of this research examines the extent to which a conditioned response to one stimulus generalizes to another. Subjects’ explicit ratings of negativity or anxiety do not seem to generalize across stimuli, while involuntary startle responses do generalize (Andreatta et al. 2015). Subjects that are not explicitly aware of the relationship between conditioned and unconditioned (painful) stimuli have generalized electrodermal responses to the experimental context up to one month later, and exhibit more avoidance behavior (Grillon 2002). Though generalization is more pronounced in those diagnosed with anxiety and other psychiatric disorders (Lissek et al. 2014; Tinoco-Gonzalez et al. 2015), this also occurs in healthy subjects (Ahrens et al. 2016). Assimilating these findings broadly, past experience with pain may lead to implicit memories that affect behavior, and awareness of pain context may affect the emergence of maladaptive responses, particularly in those with psychiatric disorders.

We examined possible measures of implicit memory as secondary outcomes. The first measure was repetition priming, that is change in average response time during the encoding task. Decreased response time with repeated exposure to the same stimuli is an expected effect of practice, but we hypothesized that pain-paired words would show a blunting of this facilitation. This prediction was based on the expectation that concomitant pain stimulation would distract attention and impair the priming effect that would otherwise shorten response time when making a different decision about a word heard previously in the experiment. We also monitored electrodermal activity and heart rate, to assess for sympathetic changes seen with repeated exposure to words during both the encoding task and during memory testing (when no pain stimuli are employed). We hypothesized that these physiologic responses would not show generalization; only pain-paired words were expected to trigger a sympathetic response, as subjects were anticipated to be explicitly aware of the pain-pairings.

### Subjects

Data were acquired from healthy volunteer subjects between the ages of 18 and 30 who were recruited from the university community. They received either course credit or $10 per hour as compensation. Eligibility was determined by self-report of exclusionary criteria and also by performance on a brief memory task. All subjects were free from significant memory impairment, hearing loss, sleep apnea, chronic pain, other chronic medical problems, neurologic and psychiatric diseases, as well as the use of antidepressants, antipsychotics, antihistamines, antianxiety medication, stimulants, sleep aids, and pain medication. The study was approved by the University of Pittsburgh Institutional Review Board (PRO16110197) and conforms to all relevant standards for the ethical and responsible conduct of research.

### Procedure

#### Screening Procedures

Subjects came into the testing location for two visits separated by at least one day. The first visit involved a paper and pencil screening to determine eligibility and to obtain responses to a series of psychometric questionnaires, including an assessment of memory ability. The brief memory task consisted of a series of 30 word pairs printed on a sheet. Subjects marked whether they thought the words in each pair were related or unrelated. Subjects then completed four psychometric questionnaires: State-Trait Inventory for Cognitive and Somatic Anxiety (STICA) (Gros et al. 2007), Pain Anxiety Symptom Scale - Short Form 20 (PASS) (McCracken and Dhingra 2002), Pain Vigilance and Awareness Questionnaire (PVAQ) (Roelofs et al. 2003), and Pain Catastrophizing Scale (PCS) (Osman et al. 1997). After spending several minutes on these questionnaires, subjects did a memory test in which all words seen previously were printed, with half of the words paired differently. Subjects were asked to indicate whether the words had previously been seen together as a pair or were re-paired.

#### Learning Task

On arrival for the second visit, subjects completed the study consent process. They were then oriented to the experiment and the nerve stimulator. The amplitude of stimulation was titrated to a targeted subjective rating of 7 out of 10 pain, using a numerical rating scale in which 0 was anchored to no pain and 10 to the worst pain imaginable. Prior to the learning portion, subjects went through a brief practice session in which they heard six words and made each of the six categorization decisions used subsequently in the experiment. One of the words in the practice session was heard twice and was followed each time by a 1.5-second electric shock. After the practice session, a pain rating was again obtained, and the nerve stimulator intensity adjusted if necessary to achieve a 7/10 rating. After this adjustment, the intensity of the stimulation was not further manipulated during the subsequent experiment, and the current flow (which was maintained at a constant value for all stimulations) was recorded. During the learning portion, subjects heard words through headphones and made decisions about them while seated at and interacting with a laptop computer. The learning portion had two conditions: the Pain condition during which every other word was paired with a painful electric shock and the No Pain condition, during which no pain was experienced. When shocks did occur, the onset was immediately after the end of the word and it was 1.5 seconds in duration. Subjects were able to respond any time after the start of the word being played, as well as during the subsequent shock, if one occurred. The maximum response time (RT) window was six seconds, after which the next word would be presented automatically. Each subject’s retrospective pain rating was obtained immediately following the Pain condition. Each of the two conditions had 50 different words that were repeated three times, with randomized word order. Within each repetition, subjects made one of six categorization decisions about the word: 1) whether it moves or not, 2) whether it is living or not, 3) whether it is natural or not, 4) whether it would fit in a shoebox or not, 5) whether it is a place or event or not, and 6) whether it is typically used or not. Subjects responded by pressing a key with their index finger to indicate a “yes” response or with their middle finger to indicate a “no” response.

Words in the pool were such that approximately 50% had yes and no responses for each categorization question. However, this constrained the moves, living, and natural decisions to be grouped together and used in one condition, while the shoebox, place, and use decisions were used in the other. The order of the Pain vs. No Pain conditions, the assignment of a set of decisions to a condition, the order of the decisions within a condition, and the order of the words within a trial were randomized for each subject with the constraint that half of the subjects received the pain context first.

#### Memory Testing

Explicit memory testing occurred after the learning portion, following a break of several minutes for instructions. Recognition testing was employed, using the Remember-Know-New (RKN) scheme (Migo et al. 2012). Subjects were given a printed sheet describing the Remember, Know, and New responses for the RKN procedure and were asked to read it. These instructions parallel those established by previous investigations (Rajaram 1993), and a copy is available as an Appendix to the supplementary materials. Subjects were asked to explain the RKN procedure back to the investigator, to ensure understanding of the differences between responses. A standardized recording summarizing these instructions was also played to the subject before beginning the RKN test. All the words previously heard in the learning portion of the experiment were then played, as well as an equal number of foils. The order of the words was randomized, and the assignment of a word from the word pool to be a foil versus used in the experiment was also random. Physiologic monitoring was continued during the RKN portion of the experiment, but no shocks were delivered.

### Stimuli

Auditory stimuli were randomly drawn from a bank created for this experiment which contained 720 English nouns. Words were obtained from the English Lexicon Project (Balota et al. 2007) database (http://elexicon.wustl.edu/). The common logarithm of the Hyperspace Analogue to Language (HAL) frequency was used to narrow words, using log(HAL) greater than 6 and less than or equal to 11 as inclusion criteria. Words were further constrained to contain between 1 and 4 syllables and between 4 and 8 phonemes. Words were then reviewed by one of the investigators to prune this list. Exclusion criteria included words that were: proper nouns, the plural form of a word already included, vulgar or otherwise objectionable, pronounced similarly to a word already included, ambiguous based on pronunciation (e.g. red vs. read), Greek letters, a slang or colloquial term, representative of an abbreviation, or starting with the same sub-word as a word already included.

Digital recordings were made by one male member of the study team speaking each word. Recordings were captured and edited using the open source software, Audacity 2.1.0 (http://audacity.sourceforge.net/) and saved as mp3 files. For timing alignment, all sound files were adjusted so that the length was exactly 750 milliseconds (ms) and the end of the word coincided with the end of the file.

### Equipment

Pain stimuli were generated using an electric nerve stimulator (EZstim II, Life Tech, Inc, Norcross, GA). Two electrodes were attached to the lateral aspect of the subject’s left index finger, straddling the proximal interphalangeal joint. Subjects were then connected to physiologic monitoring equipment by applying common electrodes to their chest for electrocardiogram monitoring, though these data are not reported here. Electrodermal activity (EDA) was acquired from the left palm, with electrodes on the hypothenar and thenar eminences. The left palm was cleaned with distilled water and allowed to air dry prior to electrode application (EDA Isotonic Gel Electrodes, Item EL507, BIOPAC Systems, Goleta, CA). Electrodes were also prepared by adding a small amount of Isotonic Recording Electrode Gel (Gel 101, BIOPAC Systems, Goleta, CA). Data were digitized using a BIOPAC MP160 (BIOPAC Systems, Goleta, CA) data acquisition unit and Acqknowledge version 5.0 (BIOPAC Systems, Goleta, CA), running on a Windows 10 laptop PC.

All parts of the experiment were implemented with E-Prime version 2.0 (Psychology Software Tools, Sharpsburg, PA). The pairing of words with electric shocks was accomplished using custom hardware that allowed E-Prime to control relays via a serial-emulated USB connection. The nerve stimulator was modified so that the push-button switches on the front of the device could be closed electronically using computer control. Response time data were logged by E-Prime during the experiment, and these intervals were synchronized to begin at the onset of the word stimulus. A 5 V square-wave logic signal indicating the timing and type of each word being played was generated by a separate bank of relays, also controlled by E-Prime. These trigger signal waveforms were recorded using the BIOPAC and allowed for alignment of the experimental events with physiologic data recordings.

### Data Analysis

Signal Detection Theory was used to estimate memory sensitivity, separate from response bias. To score memory screening results, hits were correct pair judgments, and false alarms were pairs incorrectly identified as previously-paired. A cutoff of d′ > 0.5 on the memory screen was used to determine eligibility for the second part of the experiment and no subjects were excluded based on memory performance. Memory performance for the main experiment was calculated similarly.

All statistical analysis was carried out in SPSS Statistics 23 using 95% confidence interval (CI) limits (alpha = .05) to determine statistical significance. Histograms were generated with SPSS. Other data were organized, sorted, and displayed using Microsoft Excel 2016. Outlier removal was performed using RStudio (version 1.0.153, https://www.rstudio.com/) running R version 3.2.5. The Median Absolute Deviation (MAD) was calculated as the absolute value of the difference between the median and each RT value (Leys et al. 2013). Data with MAD > 3.5 were identified as outliers. Incorrect RKN responses were removed from RT analysis. For learning response times, a repeated measures general linear model analysis was run with a Bonferroni adjustment and carried out on log-transformed data. Transformed RKN response times were analyzed using a paired samples t-test, comparing Remember and Know responses within each condition separately.

Memory testing data were assessed by calculating d′ (as described for the screening memory test) for the RKN responses. The hit rates for Remember and Know responses were calculated individually as the proportion of previously heard words designated with those responses by the subject. The false alarm rate was calculated as the proportion of foils designated with either a Remember or Know response. The d′ score thus reports the subject’s ability to discern true old items from the background of new items, effectively accounting for individual response bias.

## Results

### Subject Demographics and Pain Ratings

From the screened subjects meeting the entry criteria, 31 subjects went on to participate in the full experiment. One subject (number 12) was excluded due to a technical error. Data presented is from a cohort of 30 subjects (22 female) with age in years 23.3 ± 2.8 (mean ± standard deviation). Nerve stimulator intensity in mA was 9.9 ± 3.3. Actual pain ratings obtained just after the practice session were 6.5 ± .7 out of 10. Pain ratings following the Pain condition were 6.2 ± 1.1 out of 10.

### Psychometric Testing

Psychometric testing data were used in exploratory fashion to predict memory performance results, as in previous work (Forkmann et al. 2013; Forkmann et al. 2016). Plots of the STICA, PVAQ, PASS, and PCS against memory performance for each subject are shown in Supplementary Figures 3 and 4. Surprisingly, scores from these healthy pain-free young adults spanned the scale for all four psychometric tests, indicating a wide variety of preexisting experience with pain and anxiety. The considerable spread in psychometric data in this relatively small cohort of subjects precludes determining any significant relationships. Thus, no differences between experimental conditions based on prior measures of anxiety and pain attitudes can be reliably concluded with our data, and these are presented as exploratory results only.

### Response Time

Not surprisingly, all RT data were skewed right. Several transformations were explored, and the common logarithm (log_10_) was found to provide an approximately normal distribution of values. Histograms are shown in Supplementary Figures 5 and 6. Response time data are presented with outliers removed (technique described in the methods). For the RT data, outliers represented 3.7% of learning period responses and 4.3% of RKN responses.

Response times in ms for the three pain-pairing conditions across the three repetitions in the learning trials are shown in Figure 2. The RTs (in ms) for pain-paired words during trial 1 (mean = 1469, 95% CI = 1355-1596) were significantly greater than trial 2 (P = .001, mean = 1343, 95% CI = 1247-1442) as well as trial 3 (P < .001, mean = 1318, 95% CI = 1247-1442). For words in the No Pain Mixed condition, RTs in trial 1 (mean = 1517, 95% CI = 1400-1648) were greater than those in trial 2 (P = .001, mean = 1396, 95% CI = 1306-1493) and trial 3 (P = .003, mean = 1390, 95% CI = 1276-1517). The same pattern was observed for the No Pain Alone condition, where RTs from trial 1 (mean = 1435, 95% CI = 1343-1535) were significantly greater than trial 2 (P = .001, mean = 1327, 95% CI = 1242-1419) and trial 3 (P = .014, mean = 1327, 95% CI = 1213-1452). In comparing the three conditions collapsed across all three trials, the No Pain Mixed condition had significantly greater responses times (mean = 1432, 95% CI =1330-1542), compared to the No Pain Alone condition (P = .007, mean = 1361, 95% CI = 1268-1459).

**Figure 1.**
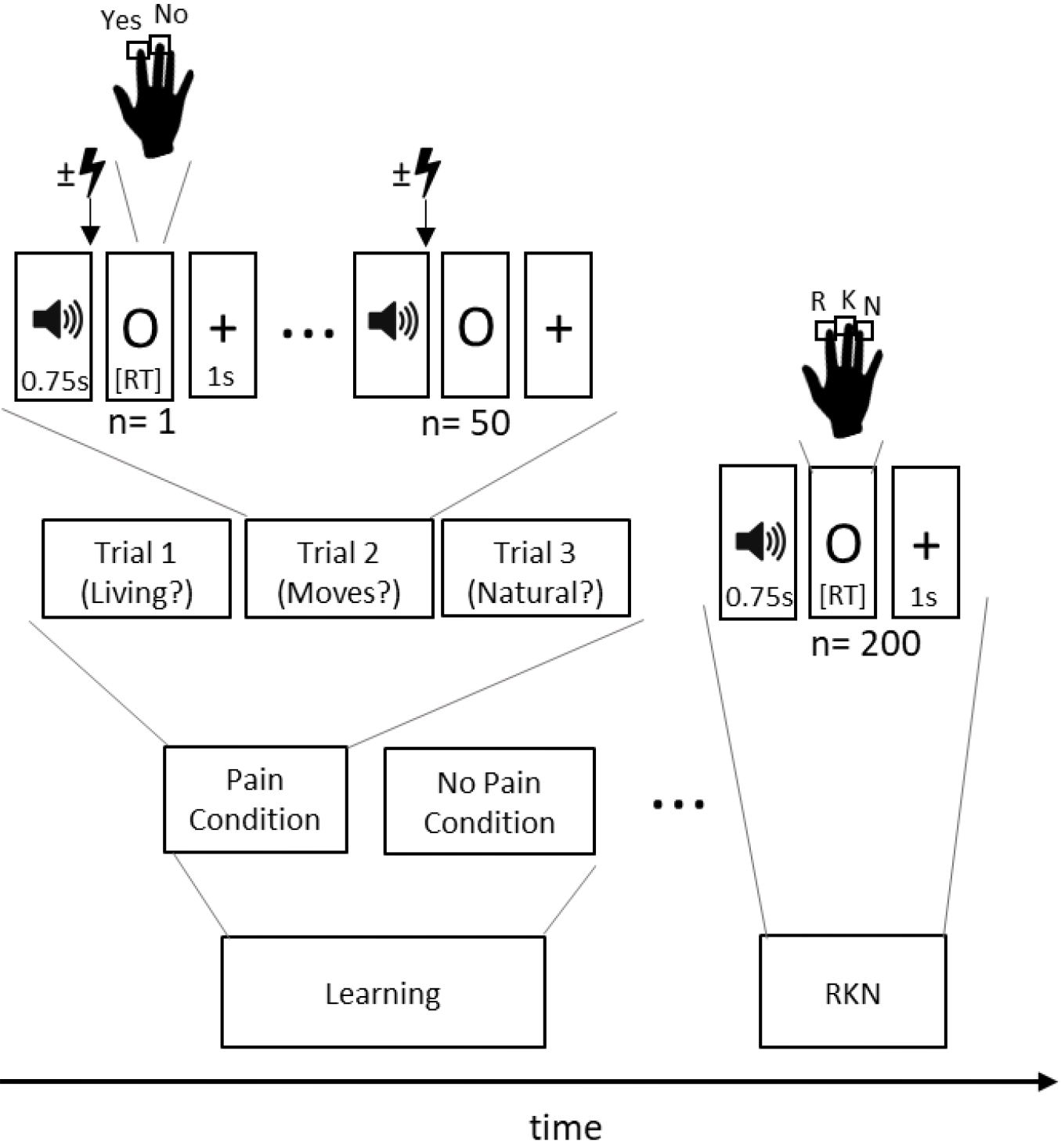
Graphical depiction of experimental design. The two main portions of the experiment are shown as blocks at the bottom. Within the different conditions (randomized order) were 3 trials using the same 50 words. Every other word received a painful electric nerve stimulation in the Pain Condition, which occurred immediately after the word finished playing.

**Figure 2.**
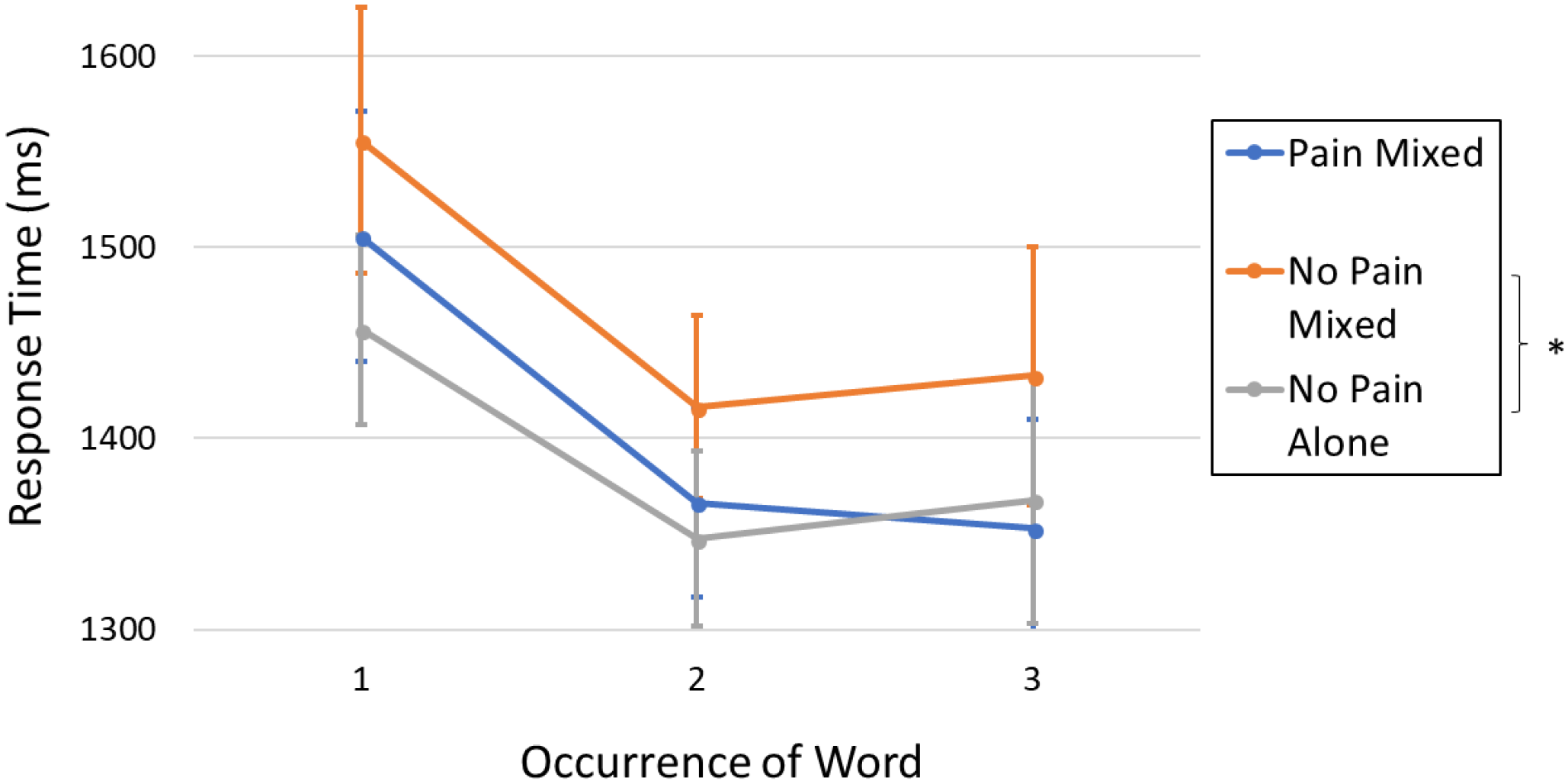
Response times over the three learning trials for each word-type. Significant differences between occurrence and between word-types are indicated with an asterisk (*).

The RKN test RT data are shown in Figure 3. As expected, Know response times were significantly longer than the Remember responses within each word-type (No Pain Mixed, P = .014, No Pain Alone, P < .001, and Pain Mixed, P = .002). Know responses for all three word-types were also significantly longer than New response times (P < .001). When comparing within Remember and Know responses individually, there were no significant differences between the three different word-types. Similarly, RTs for new words were not significantly different between any of the three word-types when collapsing across Remember and Know responses.

**Figure 3.**
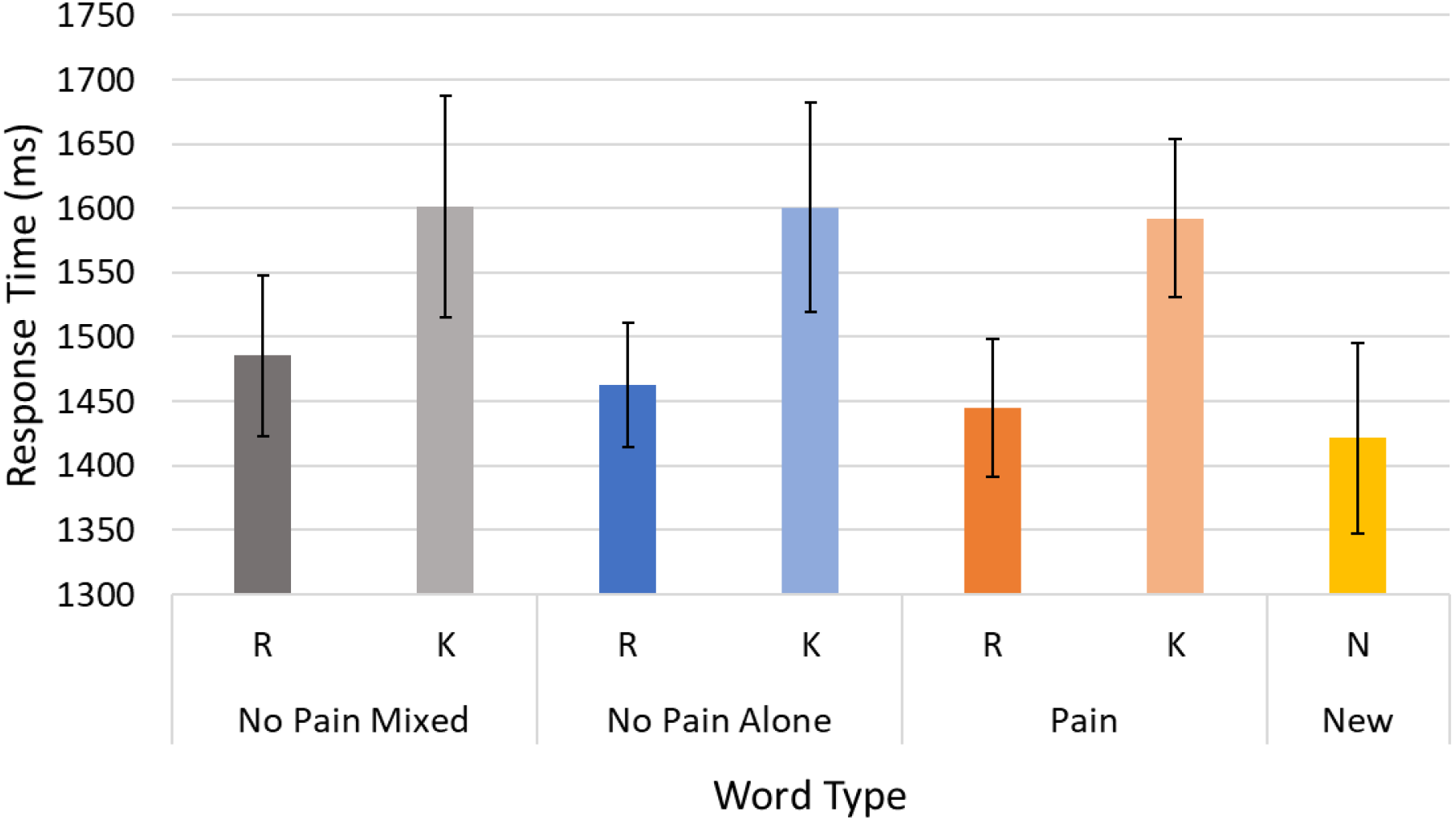
Response times for Remember (R), Know (K), and New (N) responses during the RKN testing portion, including only correct responses. R and K responses were significantly different within each word-type, and K responses for each word-type were all significantly different from N responses.

### Memory Testing

The d′ values calculated for Remember and Know responses are shown in Figure 4. Panel A compares memory performance within the Pain condition, and there were no significant differences between Pain Mixed and No Pain Mixed words. As shown in Panel B, there was a significant effect of pain context, with No Pain Alone words having better recognition than No Pain Mixed words. Specifically, Remember d′ was significantly higher (P = .028) in the No Pain Alone condition (mean = 2.68, 95% CI = 2.38-2.99) compared to the No Pain Mixed condition (mean = 2.42 95% CI = 2.12-2.74). The opposite pattern was observed for Know responses, where the No Pain Mixed condition (mean = 1.50, 95% CI = 1.15 - 1.86) exhibited significantly higher d’ values (P = .003) than the No Pain Alone condition (mean = 1.28, 95% CI = .92 - 1.64).

**Figure 4.**
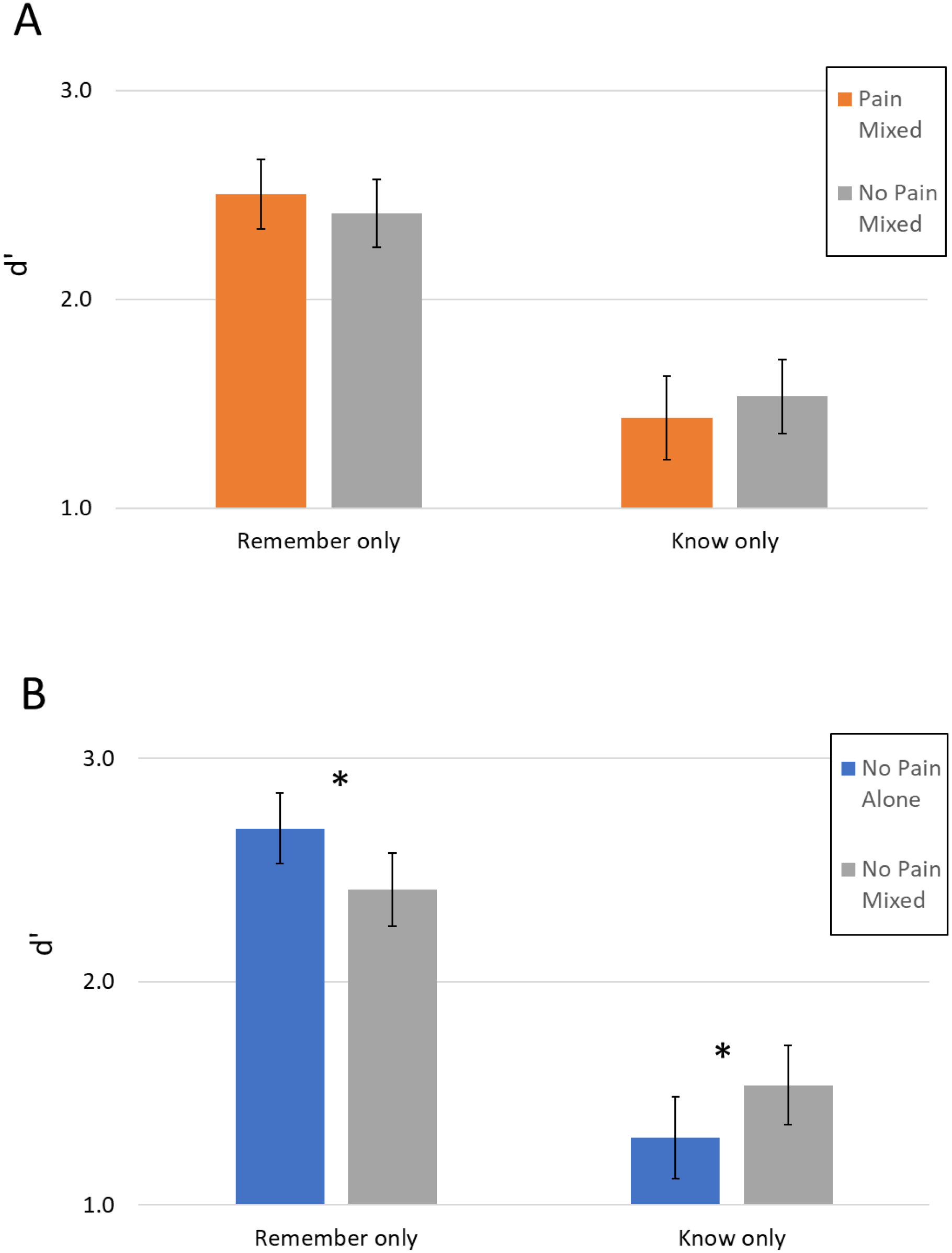
Average d′ values for Remember and Know responses for each word-type. Panel A compares the Pain Mixed and No Pain Mixed words. Panel B compares the No Pain Alone to No Pain Mixed words. Error bars display standard error. Significant differences are indicated with an asterisk (*).

The percentage of Remember responses out of total hits is shown in Figure 5. Although this measure does not account for false positive responses as d′ does, the rates of false positive responses were not significantly different across experimental word-type. Thus, Figure 5 graphically shows the proportion of responses indicating recollection (Remember) versus familiarity (Know). This proportion was significantly higher in the No Pain Alone condition (mean = .65, 95% CI = .56 - .74), compared to the No Pain Mixed condition (mean = .57, 95% CI = .47 - .67, P = .016). Notably, when asked at the end of the experiment, no subject could explicitly remember more than four words that were paired with pain, suggesting that they were not explicitly aware of the association between conditioned and unconditioned stimuli.

**Figure 5.**
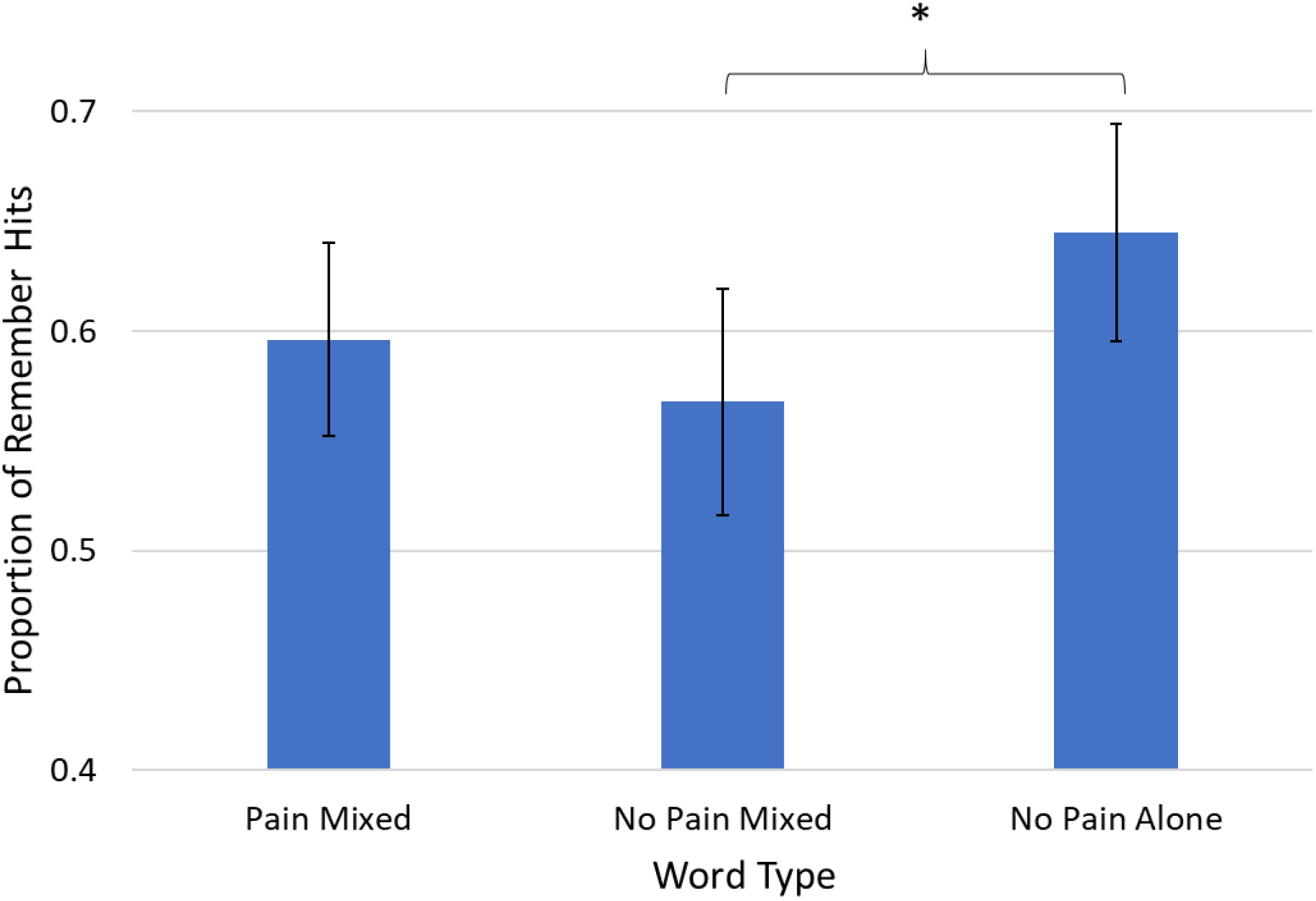
Proportion of Remember hits out of total hits for each word-type. Error bars display standard error. Significant differences are indicated with an asterisk (*).

### Electrodermal Activity

EDA results were tabulated separately for the Pain Mixed and No Pain Mixed conditions, but there were very few EDA responses for the No Pain words. Therefore, only the results from the Pain Mixed condition were analyzed. Most notably, during the RKN testing, less than 10% of all words elicited an EDA response, and the percentage of words with EDA responses did not significantly differ between conditions (P > 0.05 for all comparisons). Results for the EDA responses seen during the learning portion of the experiment are presented in the Supplementary Materials.

## Discussion

### Response Times During Learning Portion

We anticipated a decrease in RT with repeated word occurrence, particularly comparing the first to the subsequent two occurrences. This significant decrease over time, demonstrated in Figure 2 is due in part to a non-specific effect of practice of the categorization decision task, as well as a benefit of repetition of the same words (though with different decisions). The RT plateau between the second and third trials suggest that this practice effect is maximized after the first repetition. However, we also demonstrate a significant difference in response times between the two no pain conditions. Averaged across all three occurrences, No Pain Mixed words had slower responses than the No Pain Alone condition. This demonstrates an interruptive effect for pain, which seemed to slow subject responses to interleaved nonpain items (No Pain Mixed items). Though interesting, this RT decrease is not specifically indicative of an implicit memory effect. A better demonstration of a priming effect, which could be shown by a significant interaction between occurrence number and word-type, was not found in our data.

It is worth pointing out that subjects could have intentionally spaced out the occurrence of shocks by delaying their responses to either or both word-types during the Pain and No Pain Mixed conditions. Though shocks immediately followed the pain-paired items, the interval between items (and thus between the one shock and the next) was extended by the subject’s response time if it exceeded 1.5 seconds. Thus, intentional or subconscious delays in responding could have affected the Learning RT results and potentially interfered with memory encoding by cognitive distraction. However, the relatively short response intervals that were obtained (see Figure 2 and, for more detail, Supplementary Figure 5) argue against this being an issue. Subjects were also debriefed generally about their experience in the experiment, and none reported consciously employing such a strategy.

### Explicit Memory Testing

It may be worthwhile to distinguish our work from the large body of literature examining pain’s effect on *working* memory (Berryman et al. 2013). It is accepted that working memory and long-term memory recruit distinct cerebral resources, with working memory being an attention-demanding cognitive task involving prefrontal cortex (D’Esposito 2007) while long-term memory encoding is dependent on medial temporal lobe memory structures (Wais 2008). Though encoding into the working memory buffer is a necessary step prior to long-term storage, poorer performance on a task assessing working memory performance can be inversely correlated to incidental encoding into long term (Zanto et al. 2016).

In the present study, an item-specific effect of pain on explicit memory was not demonstrated. This may be explained, in part by the alternating design with respect to shocks. Working memory resources, critical to bind items into long-term memory, can be consumed by the prior experimental events, affecting performance on the subsequent item (Popov and Reder 2018). However, pain context did affect memory, as stronger explicit memory results were seen for No Pain Alone words, compared to No Pain Mixed words. There was also a corresponding shift to more familiarity judgements for No Pain Mixed words, with significantly greater d′ for Know responses, compared to No Pain Alone words. Figure 5 illustrates that these differences are driven by a shift in responses from Remember to Know, as the percentage of total hits with a Remember response is significantly lower for No Pain Mixed words, compared to No Pain Alone words. Pain Mixed words demonstrated intermediate average d′ values for both Remember and Know responses, but differences between the Pain Mixed words and the two types of non-pain words were not statistically significant. Bearing in mind the potential for a tradeoff between speed versus accuracy during the RKN testing, it is important to note that the RKN response times (Figure 3) are not demonstrably different across word-types. This suggests that differences in d′ are directly reflective of strength of memory encoding. Taken together, the RKN results indicate that experimental pain context impaired explicit memory formation. An interesting aspect of this effect is that the decrease in recollection, which we expected for pain-paired items, was surprisingly more pronounced for the non-pain items that preceded and followed the pain-paired words.

The expectation of coming noxious stimuli can affect the impact that pain has on performance for adjacent cognitive tasks that are not directly associated with pain. Unpredictability of an experimental pain stimulus is associated with higher ratings of anxiety, negative valence, and intensity (Carlsson et al. 2006). Previous studies have suggested that a warning cue preceding painful shocks had no effect on explicit memory performance (Forkmann et al. 2016). However, in a study in which subjects received a cue indicating the frequency of shock would be 20, 50, or 80 percent, a parabolic relationship was demonstrated, with highest memory for items in the 50% probability condition (Bauch et al. 2014). In the present study, the schedule for pain-pairing was consistent and described to subjects in advance, such that the shocks associated with pain-paired words were completely predictable. However, it is possible that the anticipation of a coming shock had a distracting effect for preceding words, which would be the No Pain Mixed words. Subjects might then experience a partial release of this anxiety when the shock did occur. That is, during the Pain condition of the experiment, the longest time interval in which no shock will occur starts immediately after a shock is received, including the latter portion of the decision-making period before responding to the pain-paired word. A revised experiment with pseudorandom timing of pain stimuli is currently underway to further explore the effect for pain-paired experimental events on memory of proximate non-pain items.

Previous studies examining the effect of acute pain on long-term memory have demonstrated different effects, likely due to other variations in experimental design. One factor that may influence performance is subject expectation for the effect of pain on memory. Subjects’ preconception that pain would impair memory (rated on a numerical scale) was correlated to decreases in both recollection and familiarity (Forkmann et al. 2016). Secondly, timing of the experimental pain stimulus relative to incidentally encoded cues may have an impact on the results. Studies with the painful stimulus coincident with the experimental stimulus have shown an interruptive effect for pain on memory (Forkmann et al. 2013) while pain enhanced memory when it followed neutral visual scenes by short delay of 50 ms (Schwarze et al. 2012). Notably, this enhancement was only seen in a cohort of subjects with 24-hr follow-up testing, compared to a group with testing a few minutes after the encoding portion of the experiment (Schwarze et al. 2012). This suggests that there could be an effect of memory enhancement when allowing time for memory consolidation. However, a second cohort had next-day memory testing, and, except for being in an MRI scanner, they performed the same experiment but showed no difference in memory performance (Schwarze et al. 2012). This raises the possibility of fragility of the results as an explanation of pain’s variable effect on long-term memory

Another notable difference between our study and those demonstrating a pain-related enhancement effect on memory (Schwarze et al. 2012) is the difference in latency period length prior to memory testing.

While previous studies had a latency period of one day, explicit memory testing in the present study occurred a few minutes after the final part of the encoding experiment. This timeframe and the total number of experimental items being tested (100) exceed the capacity for working memory, such that we are exclusively interrogating long-term memory. However, our design does not allow for any potential consolidation processes to occur. We have another study currently underway to examine the effect that a relatively longer follow-up period would have on memory, as longer time for consolidation in a similar paradigm may show a different result.

Putting our explicit memory results in the context of the broader literature, one could interpret our equivocal differences specifically for pain-paired items, compared to both types of non-pain items, as a null result. However, the stronger recollection performance for No Pain Alone versus No Pain Mixed words (neither of which are directly associated with painful stimulation) demonstrate that pain context impairs recognition memory. This suggests that, at least in a 1:2 pain-pairing paradigm, this contextual effect predominates over an item-specific effect for explicit memory. That is, pain’s generalized effect on recollection predominates over the effect pain has on explicit memory for specific pain-paired items.

## Conclusions

For aurally delivered words about which subjects performed a categorization decision, experimental painful stimulation affected memory performance. Explicit memory, indicated by Remember-Know testing, was most affected. Compared to non-pain words heard in a pain-free portion of the experiment, non-pain words heard in the context of other pain-associated words were not as deeply encoded. This was demonstrated by less recollection and a compensatory increase in familiarity responses. An important accompanying finding is that response times to repeated experimental items were slower for non-pain words heard in the context of pain-paired words, compared to non-pain items heard in a pain-free context. Taken together, these results suggest that, compared to a pain-free context, acute pain stimulations create an experiential context wherein memory formation is affected, even for items not directly followed by painful stimulations.

## Acknowledgements

This work was supported by a seed grant from the Department of Anesthesiology, University of Pittsburgh, School of Medicine. Further support was provided by a Mentored Research Training Grant (to KMV) from the Foundation for Anesthesia Education and Research (FAER). Salary support for KMV during the initial phase of this project came from an institutional training grant from the National Institutes of Health (T32GM075770). Subject recruitment was assisted by the University of Pittsburgh Clinical Translational Science Institute Research Participant Registry, a project supported by the National Institutes of Health through grant number UL1TR000005.

## References

Ahrens LM, Pauli P, Reif A, Muhlberger A, Langs G, Aalderink T, Wieser MJ. 2016. Fear conditioning and stimulus generalization in patients with social anxiety disorder. J Anxiety Disord 44: 36–46.

Andreatta M, Leombruni E, Glotzbach-Schoon E, Pauli P, Muhlberger A. 2015. Generalization of Contextual Fear in Humans. Behav Ther 46: 583–596.

Bauch EM, Rausch VH, Bunzeck N. 2014. Pain anticipation recruits the mesolimbic system and differentially modulates subsequent recognition memory. Hum Brain Mapp 35: 4594–4606.

Berryman C, Stanton TR, Jane Bowering K, Tabor A, McFarlane A, Lorimer Moseley G. 2013. Evidence for working memory deficits in chronic pain: a systematic review and meta-analysis. Pain 154: 1181–1196.

Carlsson K, Andersson J, Petrovic P, Petersson KM, Ohman A, Ingvar M. 2006. Predictability modulates the affective and sensory-discriminative neural processing of pain. NeuroImage 32: 1804–1814.

Castegnetti G, Tzovara A, Staib M, Paulus PC, Hofer N, Bach DR. 2016. Modeling fear-conditioned bradycardia in humans. Psychophysiology 53: 930–939.

D’Esposito M. 2007. From cognitive to neural models of working memory. Philos Trans R Soc Lond B Biol Sci 362: 761–772.

Forkmann K, Schmidt K, Schultz H, Sommer T, Bingel U. 2016. Experimental pain impairs recognition memory irrespective of pain predictability. Eur J Pain 20: 977–988.

Forkmann K, Wiech K, Ritter C, Sommer T, Rose M, Bingel U. 2013. Pain-specific modulation of hippocampal activity and functional connectivity during visual encoding. J Neurosci 33: 2571–2581.

Grillon C. 2002. Associative learning deficits increase symptoms of anxiety in humans. Biol Psychiatry 51: 851–858.

Gros DF, Antony MM, Simms LJ, McCabe RE. 2007. Psychometric properties of the State-Trait Inventory for Cognitive and Somatic Anxiety (STICSA): comparison to the State-Trait Anxiety Inventory (STAI). Psychological assessment 19: 369–381.

Leys C, Ley C, Klein O, Bernard P, Licata L. 2013. Detecting outliers: Do not use standard deviation around the mean, use absolute deviation around the median. Journal of Experimental Social Psychology 49: 764–766.

Lissek S, Kaczkurkin AN, Rabin S, Geraci M, Pine DS, Grillon C. 2014. Generalized anxiety disorder is associated with overgeneralization of classically conditioned fear. Biol Psychiatry 75: 909–915.

McCracken LM, Dhingra L. 2002. A short version of the Pain Anxiety Symptoms Scale (PASS-20): preliminary development and validity. Pain Res Manag 7: 45–50.

Migo EM, Mayes AR, Montaldi D. 2012. Measuring recollection and familiarity: Improving the remember/know procedure. Consciousness and cognition 21: 1435–1455.

Osman A, Barrios FX, Kopper BA, Hauptmann W, Jones J, O’Neill E. 1997. Factor structure, reliability, and validity of the Pain Catastrophizing Scale. J Behav Med 20: 589–605.

Popov V, Reder LM. 2018. Frequency Effects on Memory: A Resource-Limited Theory.

Rajaram S. 1993. Remembering and knowing: two means of access to the personal past. Mem Cognit 21: 89–102.

Roelofs J, Peters ML, McCracken L, Vlaeyen JW. 2003. The pain vigilance and awareness questionnaire (PVAQ): further psychometric evaluation in fibromyalgia and other chronic pain syndromes. Pain 101: 299–306.

Schiller D, Monfils MH, Raio CM, Johnson DC, Ledoux JE, Phelps EA. 2010. Preventing the return of fear in humans using reconsolidation update mechanisms. Nature 463: 49–53.

Schwarze U, Bingel U, Sommer T. 2012. Event-related nociceptive arousal enhances memory consolidation for neutral scenes. J Neurosci 32: 1481–1487.

Tinoco-Gonzalez D, Fullana MA, Torrents-Rodas D, Bonillo A, Vervliet B, Blasco MJ, Farre M, Torrubia R. 2015. Conditioned Fear Acquisition and Generalization in Generalized Anxiety Disorder. Behav Ther 46: 627–639.

Wais PE. 2008. FMRI signals associated with memory strength in the medial temporal lobes: a meta-analysis. Neuropsychologia 46: 3185–3196.

Zanto TP, Clapp WC, Rubens MT, Karlsson J, Gazzaley A. 2016. Expectations of Task Demands Dissociate Working Memory and Long-Term Memory Systems. Cereb Cortex 26: 1176–1186.

